## Supplementary Materials for "Modelling The Effect of MUC1 on Influenza Virus Infection Kinetics and Macrophage Dynamics"

### Supplementary Material 1

#### Model parameters and Priors

Table 1 gives the prior distribution for estimated parameters. Note that we estimate $\beta$, $p$, $\kappa_M$, $s$ and $\phi$ in logarithmic space. For example, the estimated ranges of $log_{10}(\beta)$ were (-8, -4) from literature, which indicate the estimated ranges of $\beta$ were $(10^{-8}, 10^{-4})$. We set the priors which allow us to explore a wide range of biological plausible parameter values. Table 2 gives parameter values for fixed parameters.

| Par. | Description | Estimated values from literature [Refs] | Unit | Prior |
|---|---|---|---|---|
| $\varepsilon_1$ | The effect of MUC1 on viral infectivity | - | - | Uniform(0,1) |
| $\log_{10}(\beta)$ | Rate of viral infection | (-8,-4) [1, 2] | $/([u_V]$ day) | Normal(-6,-4) |
| $\delta_I$ | Death rate of infected cells | (0.67, 4.8)[1, 2, 3] | /day | Lognormal(log(0.89),1) |
| $\log_{10}(p)$ | Viral production rate | (-6,2)[1, 2, 3] | $[u_V]$/(cell day) | Normal(-2,4) |
| $\delta_V$ | Natural death rate of virus | (4.2, 59)[1, 2, 3] | /day | Lognormal(log(28.4),1) |
| $\log_{10}(\kappa_M)$ | Phagocytosis rate of virus by macrophages | (-6,-3)[4] | /(cell day) | Normal(-6,4) |
| $\varepsilon_2$ | The effect of MUC1 on macrophage recruitment | - | - | Uniform(0,1) |
| $\delta_M$ | Decay rate of macrophages | (1/180,1/150)[4] | /day | Lognormal(log(4.2e-3),1) |
| $\log_{10}(s)$ | Supplementary rate of macrophages | (2.52, 2.63)[4] | cell/(ml day) | Normal(3,1) |
| $\log_{10}(\phi)$ | Recruitment rate of macrophages by infected cells | - | (ml cell)/cell | Normal(0,3) |

Table 1: **Priors for estimated model parameters.** $[\cdot]$ denotes the unit of variables, e.g., the unit of virus is denoted as $[u_V]$.

| Par. | Description | Values [Refs] | Unit |
|---|---|---|---|
| $g$ | Epithelial cell regrowth rate | 0.8 [5, 6] | /day |
| $\gamma_E$ | Maximal stimulation rate of naive CD8 T cells | 10 [6] | /day |
| $E_{50}$ | Half-maximal stimulating viral titer for CD8 T cells | 1e+4 | $[u_V]$ |
| $n_E$ | Number of effector T cell division cycle | 5 [6] | division |
| $\tau_E$ | Total proliferation time of CD8 T cells | 8 [6] | day |
| $\phi_E$ | Activation rate of matured CD8 T cells | 1.4e+3 [6] | /day |
| $\delta_E$ | Decay rate of CD8 T cells | 0.57 [6] | /day |
| $\kappa_E$ | Lysing rate of infected cells by CD8 T cells | 5e-5 [7] | /day/cell |
| $\gamma_B$ | Maximal stimulation rate of naive B cells | 6e-2 [6] | /day |
| $B_{50}$ | Half-maximal stimulating viral titer for B cells | 6e-2 | $[u_V]$ |
| $n_B$ | Number of B cell division cycle | 5 [6] | division |
| $\tau_B$ | Total proliferation time of B cells | 8 [6] | day |
| $\phi_P$ | Activation rate of matured plasma cells | 8 [6] | /day |
| $\delta_P$ | Decay rate of plasma cells | 0.5 [6] | /day |
| $\mu_S$ | Production rate of short-lived antibody | 12 [5, 6, 7] | $[u_A]$/cell/day |
| $\delta_{AS}$ | Decay rate of short-lived antibody | 2 [5] | /day |
| $\mu_L$ | Production rate of long-lived antibody | 4 [7] | $[u_A]$/cell/day |
| $\delta_{AL}$ | Decay rate of long-lived antibody | 0.015 | /day |
| $\kappa_{AS}$ | Neutralisation rate of virus by short-lived antibody | 0.8 [7] | $[u_A]$/day |
| $\kappa_{AL}$ | Neutralisation rate of virus by long-lived antibody | 0.8 [5, 7] | $[u_A]$/day |
| $T_{max}$ | The maximal number of epithelial cells in the upper respiratory tract | 1e+7 [1] | cell |

Table 2: **Parameter values for fixed parameters.** $[\cdot]$ denotes the unit of variables, e.g., the unit of antibody is denoted as $[u_A]$, and the unit of virus is denoted as $[u_V]$.
